# Identification of cellular-activity dynamics across large tissue volumes in the mammalian brain

**DOI:** 10.1101/132688

**Authors:** Logan Grosenick, Michael Broxton, Christina K. Kim, Conor Liston, Ben Poole, Samuel Yang, Aaron Andalman, Edward Scharff, Noy Cohen, Ofer Yizhar, Charu Ramakrishnan, Surya Ganguli, Patrick Suppes, Marc Levoy, Karl Deisseroth

## Abstract

Tracking the coordinated activity of cellular events through volumes of intact tissue is a major challenge in biology that has inspired significant technological innovation. Yet scanless measurement of the high-speed activity of individual neurons across three dimensions in scattering mammalian tissue remains an open problem. Here we develop and validate a computational imaging approach (SWIFT) that integrates high-dimensional, structured statistics with light field microscopy to allow the synchronous acquisition of single-neuron resolution activity throughout intact tissue volumes as fast as a camera can capture images (currently up to 100 Hz at full camera resolution), attaining rates needed to keep pace with emerging fast calcium and voltage sensors. We demonstrate that this large field-of-view, single-snapshot volume acquisition method—which requires only a simple and inexpensive modification to a standard fluorescence microscope—enables scanless capture of coordinated activity patterns throughout mammalian neural volumes. Further, the volumetric nature of SWIFT also allows fast in vivo imaging, motion correction, and cell identification throughout curved subcortical structures like the dorsal hippocampus, where cellular-resolution dynamics spanning hippocampal subfields can be simultaneously observed during a virtual context learning task in a behaving animal. SWIFT’s ability to rapidly and easily record from volumes of many cells across layers opens the door to widespread identification of dynamical motifs and timing dependencies among coordinated cell assemblies during adaptive, modulated, or maladaptive physiological processes in neural systems.

## Introduction

The ability to record activity from intact tissue at physiologically relevant timescales and at appropriate resolution is central to any attempt to model and understand the dynamic behavior of complex biological systems. The mammalian nervous system represents a particularly challenging example of such a system, due to the complex, three-dimensional spatiotemporal activity patterns that diverse populations of neurons exhibit as they store, retrieve, and process information at millisecond time scales across large volumes of tissue. Consequently, capturing the coordinated neural firing and statistical dependencies among cells that are thought to be important in behavior (and disease) will ultimately require simultaneous high-speed observation of neuronal timing relationships throughout large tissue volumes.

Since the introduction of synthetic (Tsien 1980), genetically-encoded Ca^2+^ (Miyawaki et al. 1997; Tian et al. 2009), and voltage sensors (Akemann et al. 2010; Kralj et al. 2011; Jin et al. 2012; St-Pierre et al. 2014; Gong et al. 2014), a number of powerful new technologies for optically recording activity signals in neurons have been introduced. Recent advances in single-cell Ca^2+^ imaging have yielded impressive results using fast three-dimensional scanning of single (Göbel, Kampa, and Helmchen 2007; Duemani Reddy et al. 2008) or multiple (Niesner et al. 2007; Cotton et al. 2013) laser beams, or by scanning a sheet of light (Tomer et al. 2012; Ahrens et al. 2013; Bouchard et al. 2015) to image many neurons by sequentially illuminating parts of a volume over time. However, such methods capture the volume (and therefore the activity of neurons across the volume) asynchronously, and are fundamentally limited by the inertial and settling constraints of the moving components. Although scanless volumetric microscopy technologies including holographic (Rosen and Brooker 2008), extended depth-of-field (Quirin et al. 2014), and multi-focus (Abrahamsson et al. 2012) microscopy exist, these methods require acquisition of multiple images for each volume, projection from three into two dimensions, or suballocation of the camera sensor for each axial plane, thereby limiting rapid three-dimensional imaging and full volumetric reconstruction. To date, no mammalian activity imaging approach has allowed synchronous full-volume acquisition with validated cellular resolution using the full scope and speed of fast cameras. Compounding this challenge, analytical tools necessary for accurate discovery of neurons in such large volumetric time series data over time remain to be developed.

In considering these challenges, we noted that certain features of light field microscopy—a method that allows the reconstruction of a full volume from a single camera frame (Marc Levoy et al. 2006; Marc Levoy, Zhang, and McDowall 2009; Broxton et al. 2013; Cohen et al. 2014)—might support synchronous volumetric imaging in mammalian preparations at rates appropriate for new generations of fast calcium and voltage sensitive dyes. Several proof-of-concept experiments have developed and demonstrated the basics of light field deconvolution (Broxton et al. 2013; Prevedel et al. 2014) and light field microscopy has been shown to work for volumetric functional imaging of neuronal activity in small, optically transparent neurobiological systems, first in a single brain region (Grosenick, Anderson, and Smith 2009) and subsequently across a zebrafish brain—all of these approaching but not fully attaining cellular resolution (Deisseroth and Schnitzer 2013; Prevedel et al. 2014; Tomer et al. 2015), and most recently across the Drosophila brain (Aimon et al. 2015). However, adapting and scaling the technology for millimeter-scale volumes in scattering mammalian preparations and achieving single-cell resolution have remained the biggest challenges.

We hypothesized that with our volume reconstruction method (Broxton et al. 2013), it might be possible to attain camera-limited imaging rates at a resolution that would allow identification and activity recording from individual neurons over significant tissue volumes in mammalian systems both in vitro and in vivo during behavior. However, beyond the volume reconstruction—already a challenging computational problem to scale to many volumes—significant analytical advances would also be required, since identification of neurons and dynamics over time in volumes consisting of ∼5-100 million voxels at rates up to 100 Hz (terabytes per data set) would require substantial development of statistical and computational methodology. In particular, machine learning methods appropriate for motion correction and cell localization in terabyte-scale data sets would be needed. To address this challenge, here we develop, validate, and apply an integrative computational imaging approach (SWIFT) enabling the synchronous acquisition and analysis of single-neuron resolution activity throughout intact scattering tissue volumes at high frame rates. Open source code implementing SWIFT will be made available online.

## Single-snapshot volumetric imaging of biological neural networks in mammalian tissue

By inserting an array of thousands of microlenses at the intermediate image plane of a standard epifluorescence microscope and positioning the camera sensor at the image plane formed by this array (Fig. 1A,B), it is possible to capture the four-dimensional light field in a single, synchronous exposure of the camera sensor (Marc Levoy et al. 2006; Marc Levoy, Zhang, and McDowall 2009). This light field image (Fig. 1D) encodes information about the entire volume under the objective, albeit at somewhat reduced lateral resolution (Supplementary Fig. 1). However, this modest reduction in lateral resolution is acceptable so long as we can still identify individual cell bodies, and worthwhile as it fundamentally enables synchronous, high-speed volumetric functional imaging. A single light field volume can be reconstructed from a single camera frame to obtain a volume by solving a large maximum likelihood problem (e.g., inverting a 5.5 million pixel by 2-100 million voxel linear system per volume; Fig. 1C), thereby estimating the most likely intensity distribution in the volume given the captured image on the sensor. Our algorithm is analogous to tomographic reconstruction and deconvolution techniques in medical imaging, but has been adapted here for single-snapshot volume fluorescence imaging (Broxton et al. 2013). We use a wave optics (i.e., physical optics) model of light transport through the microscope and the microlens array and a noise model that correctly accounts for noise sources on the image sensor (principally photon shot noise). Due to the manner in which light is focused by the microlenses, this inverse problem is well conditioned and cell-scale details can be recovered much deeper into scattering tissue than standard epifluorescence imaging allows. Fig. 1E and Supplementary Movie 1 demonstrate that such reconstruction is achievable for a complex living neural system at cell-resolution in mammalian tissue; hippocampal neurons in an acute mouse slice preparation are shown expressing the genetically-encoded Ca^2+^ sensor GCaMP6f (T.-W. Chen et al. 2013). With a 40x 0.8 NA objective, apical dendrites and the largest spines are just visible. Each full volume in a time series is reconstructed using a single camera exposure, thus the imaging rate is limited only by the available light from the sample and the maximum frame rate of the camera.

**Fig. 1.**
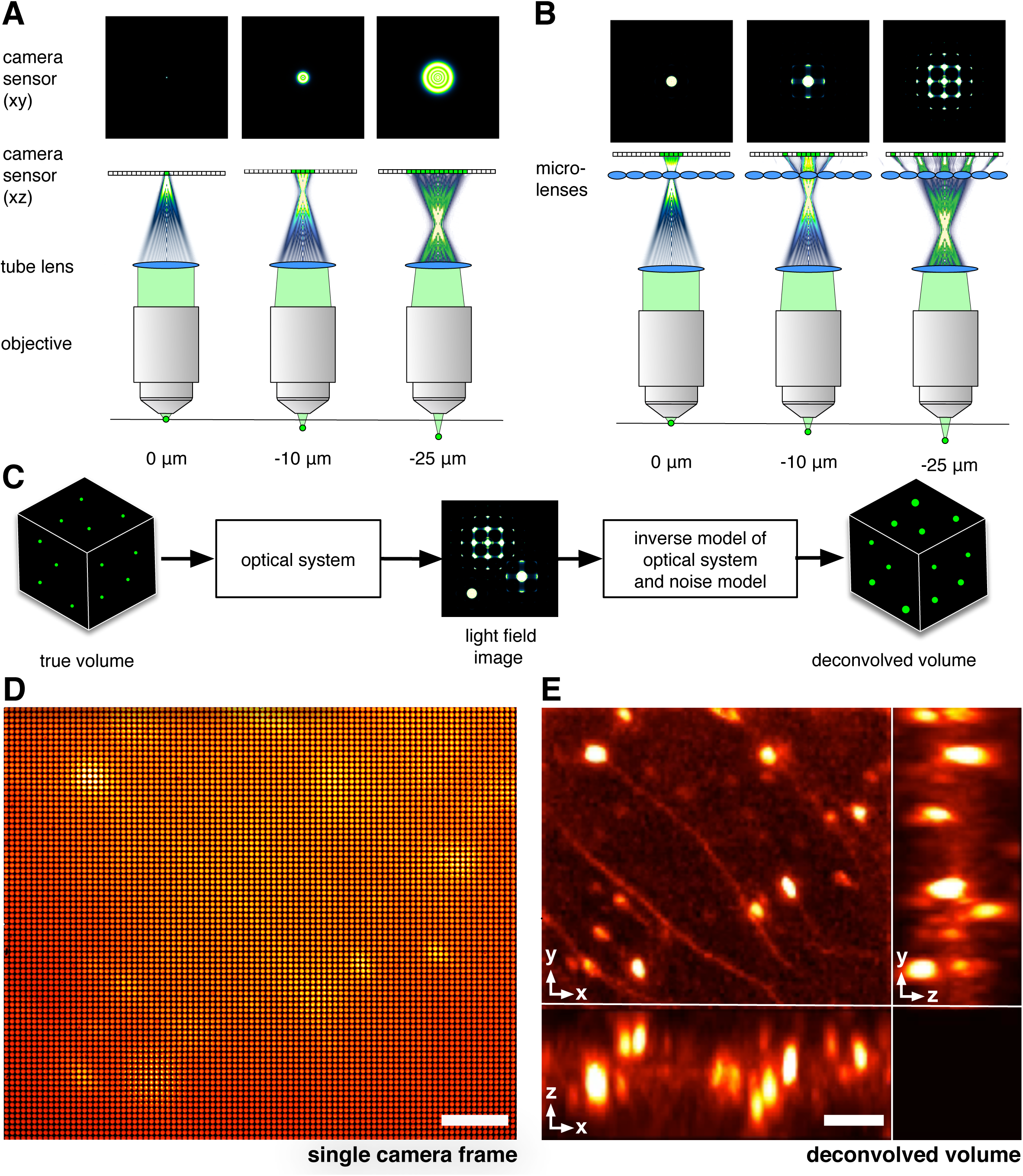
Synchronous volumetric functional imaging in mammalian tissue at cellular resolution with SWIFT. (**A**) In a conventional fluorescence microscope, a fluorescent point source moving away from the plane of focus creates an increasingly blurred image of the source, making it difficult to determine its lateral and axial position. (**B**) In a light field microscope (Marc Levoy et al. 2006), a microlens array (Methods) in the optical path refocuses this light, creating a distinctive diffraction pattern that is unique for each position of the point source in the volume. This form of optical coding enables synchronous capture of information about a full volume in a single image. (**C**) A physical (“wave”) optics description of light transport through the microscope can then be used to solve a three-dimensional deconvolution problem to reconstruct a volume from a single two-dimensional light field image (Broxton et al. 2013) (Methods). (**D**,**E**) Raw and reconstructed light field images from an acute mouse hippocampal slice expressing genetically encoded indicator GCaMP6f (T. W. Chen et al. 2013), showing this technique works even in scattering mammalian tissue. (**D**) A raw, two-dimensional light field image acquired by the camera (scale bar 330 camera pixels). (**E**) Maximum-intensity projections through the three-dimensional volume reconstructed using the two-dimensional image from **D**, showing a volume with clearly discernible neurons and apical dendrites in mouse ventral hippocampus at one instant in time (50 ms exposure, standard illumination at 475/35nm, 450 μW/mm^2^; scale bar 100 μm; volume is 200 μm deep; see also Suppl. Movie 1).

However, the rapid collection of large volume data immediately creates a problem: many tens of thousands of images recorded during a single experiment must be reconstructed into volumes, motion aligned, and then further processed to extract and analyze the spatial locations and time courses of the large numbers of neurons. To meet this challenge, we developed our own distributed data processing system that runs on the Amazon Elastic Compute Cloud (EC2) and the Google Compute Engine (GCE) cloud computing services (see Methods). Our GPU-accelerated volumetric deconvolution system is capable of reconstructing 9000 volumes per hour on 300 compute instances. Subsequent motion alignment, source extraction, and time series analysis is carried on clusters ranging from 128-1024 cores using the Apache Spark cluster computing framework (Zaharia et al. 2016) and the Thunder data analysis framework (Freeman et al. 2014).

For extraction of the locations of neural sources and their time series from motion aligned volumetric time series data, we developed new, scalable machine learning methods based on previous work in high dimensional settings (Jenatton, Obozinski, and Bach 2009; Grosenick et al. 2013; Allen, Grosenick, and Taylor 2014) to seek smooth, local spatial structures with global sparsity so that these resolved “sources” were cell-sized and cell-shaped, i.e. sparse but structured nonnegative matrix factorization. Similar methods have recently yielded state-of-the-art results on smaller problems (Pnevmatikakis et al. 2016). Along with methods to significantly accelerate computational extraction of the neural sources (Methods; runtime is about one hour for 3000 sources in a volume with two million voxels over 27000 time points), we also developed automated screening and validation procedures to increase confidence that sources used in all subsequent analyses are of neural origin (See Methods; runtime is about 10 minutes on the same data). We validated these methods and the quality of the spatial and temporal sources (see Methods). This integration of physical optics, computational optics, and statistical methods enables extraction of neural locations and activity time series simultaneously across mm-scale tissue volumes; we term this method SWIFT (Statistical Wave-optics Identification of Features in Tissue) volume imaging.

## SWIFT: fast functional recording of volumes of neurons

Standard light field deconvolution microscopy is compatible with imaging neural activity in largely transparent animals such larval zebrafish (Grosenick, Anderson, and Smith 2009; Prevedel et al. 2014; Cohen et al. 2014) and certain invertebrates such as C. elegans and Drosophila (Prevedel et al. 2014; Aimon et al. 2015), but most vertebrate neural tissues, including mammalian brains, are turbid and highly scattering (Ntziachristos 2010). However, we found that due to its light field deconvolution model SWIFT was effective for volumetric functional imaging in living mammalian neural circuitry. Beginning with acute mouse prefrontal cortical (PFC) slices virally transduced with a genetically encoded Ca^2+^ indicator (GECI) using an adeno-associated virus (AAVdj::CamKIIa::GCaMP6f; Methods) and using a 10x 0.6NA objective, we were able to simultaneously image 1.1 mm x 1.0 mm x 0.4 mm volumes spanning all cortical layers and all of prelimbic cortex (PL) in vitro with parts of cingulate cortex (Cg) and/or infralimbic cortex (IL) in a single field of view (Supplementary Fig. 2A,B; Supplementary Video 2). We found that SWIFT was in fact well suited to identifying neurons and their time series through this scattering volume of tissue even using wide-field illumination (Supplementary Fig. 2B-D; Methods). Indeed, the larger size of mouse neurons (20-30 μm soma relative to 5-10 μm soma in, for example, zebrafish fry) enabled efficient segmentation of individual cells with computationally-optimized, high-dimensional machine learning identification of smooth, local spatial structures (“structured sparsity”) (Jenatton, Obozinski, and Bach 2009; Grosenick et al. 2013) so that resolved sources of high temporal variance were the size and shape of cell bodies (Supplementary Fig. 2B). Supplementary Fig. 2C shows 12 Ca^2+^ imaging fluorescence traces with numbers corresponding to the numbered spatial filters shown in Supplementary Fig. 2B, with cells exhibiting a diversity of activity, from sparse spiking to concentrated bursts of activity similar to “UP” states previously reported in cortical slices (Cossart, Aronov, and Yuste 2003). Supplementary Fig. 2D-H further characterize the autocorrelations and signal-to-noise using whole-cell patch clamp, and by comparing 25 Hz two-photon laser scanning microscopy to 25 Hz SWIFT volume imaging in the same cells. No cross-talk was detectable between reconstructed neurons, we confirmed that the spatial sources resembled the somata of patched, back-filled cells, and found that SWIFT exhibited an approximately 2x improvement in SNR over a commercial, resonant scanning 2PSLM system (Leica SP5; See Supplementary Fig. 2 and Methods).

## Synchronous volumetric imaging across hippocampal subfields

This success of SWIFT for imaging volumes of neurons in vitro led us to explore imaging of large subcortical brain volumes in behaving mice. Following the techniques shown in previous groundbreaking work imaging in dorsal hippocampus (subfield CA1) (Dombeck et al. 2007; Barretto, Messerschmidt, and Schnitzer 2009), we implanted glass windows just above the dorsal hippocampus in C57BL/6J mice (Fig. 2A,B; Methods), moving our coordinates anterior and lateral relative to previous studies in order to allow simultaneous optical access with a 10x 0.6 NA long-working-distance water-dipping objective to parts of hippocampal subfields CA1, CA2, and CA3 virally transduced with GCaMP6f (AAVdj-CamKIIa-GCaMP6f, Methods). Starting at 2-3 weeks after viral transduction and implantation, mice could be imaged with SWIFT while head-fixed and running on an axially-fixed spherical treadmill (Fig. 2C) (Dombeck et al. 2007).

**Fig. 2.**
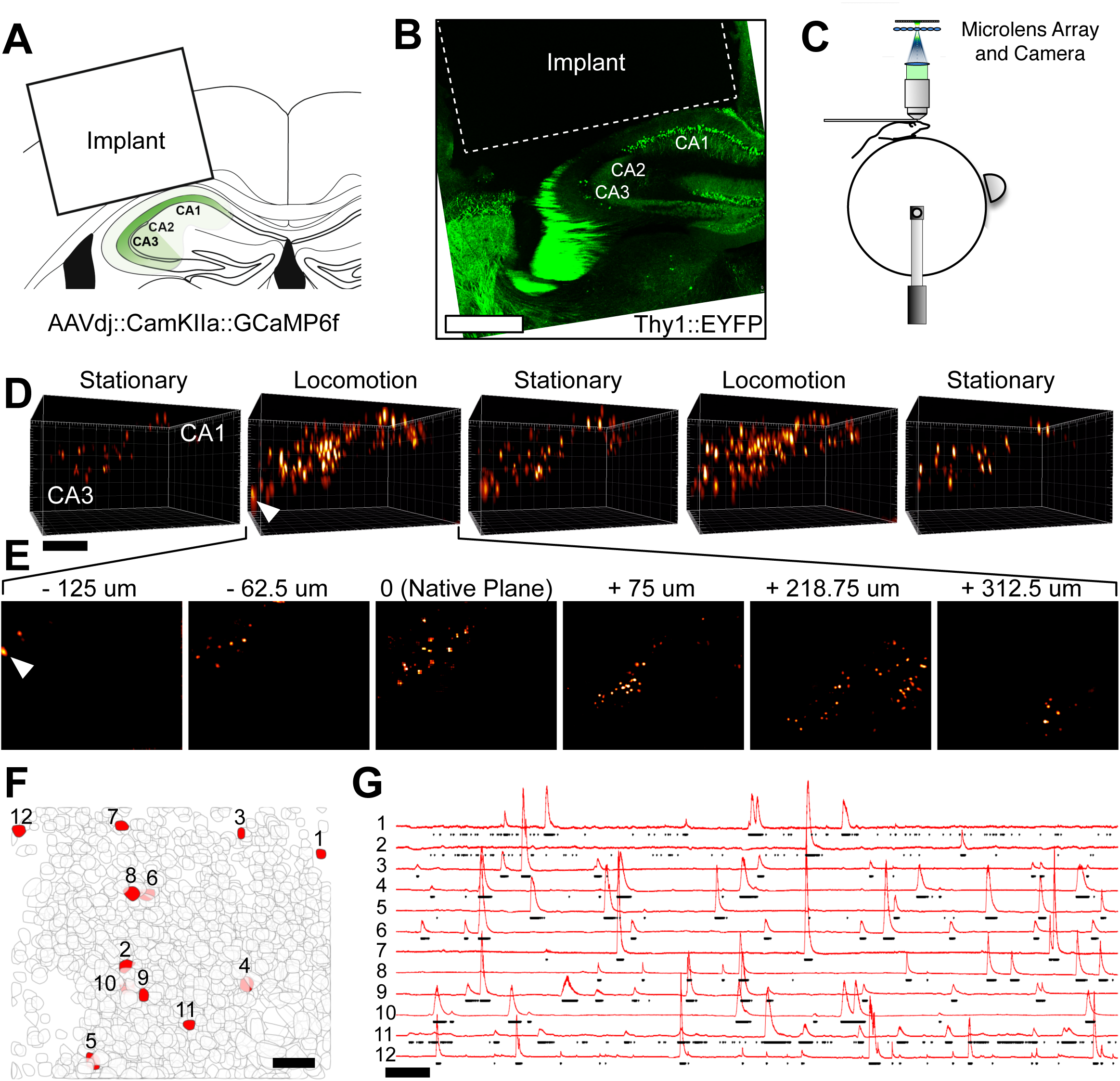
SWIFT volume imaging across hippocampal subfields in a behaving rodent. (**A**) Mice expressing genetically-encoded calcium indicator GCaMP6f in hippocampal subfields CA1, CA2, and CA3 are implanted with a cranial window allowing optical access to the dorsal hippocampus (Dombeck et al. 2007; Barretto, Messerschmidt, and Schnitzer 2009). Subfields CA1-CA3 are present in the imaging field of view. (**B**) Histology from an EYFP animal showing cannula placement and location relative to subfields. (**C**) A head-restrained mouse is imaged while running on an axially-fixed spherical treadmill. Mouse movements on the ball are measured with an optical mouse as in previous work (Dombeck et al. 2007). (**D**) Volumes from SWIFT imaging acquired synchronously at 50 Hz; each of the 5 time point volumes was taken when the mouse was either running or stationary, as labeled. The white arrow points to a neuron more than 0.5 mm below the top of the volume (scale bar 200 μm). (**E**) Z-slices through a single volume showing cells at different depths in the volume. (**F**) Active neurons across hippocampal subfields expressing genetically-encoded calcium indicator GCaMP6f and localized using a novel sparse, structured machine learning method appropriate for massive data (Methods; scale bar 100 μm). (**G**) 12 estimated calcium traces corresponding to the spatial filters of the same number in (**F**), with thresholded spike probability estimates (threshold at 5% percentile) (Vogelstein et al. 2010; Pnevmatikakis et al. 2016) (Methods, scale bar 30 sec).

Volumes were reconstructed as in the in vitro experiments above, however, in a behaving animal brain motion relative to the sensor resulted in motion artifacts between temporally adjacent volumes, as previously described (Dombeck et al. 2007). Such artifacts can significantly degrade accurate estimation of cell location and sensor transients, making image alignment a critical first step for SWIFT as for other calcium and voltage imaging modalities. However image alignment is considerably easier in SWIFT because each volume can be treated as a frame captured at a single period of time. This avoids nonlinear sampling effects known to make motion correction difficult in data acquired by scanning methods (Dombeck et al. 2007). Even so, the high dynamic range and low background signal of GCaMP6 required statistically partitioning large calcium transients from the static background fluorescence, the latter being necessary for motion correction. To accomplish this we developed a motion correction protocol using the Robust Alignment by Sparse and Low Rank (RASL) algorithm (Peng et al. 2012) (Methods), and scaled it to work on terabyte-scale data using Apache Spark (Zaharia et al. 2016). This method identifies set of three-dimensional translation and rotation corrections to each volume and allows the entire dataset to be decomposed into sparse neural dynamics superimposed upon a low-rank background signal (Peng et al. 2012). We found that RASL outperformed normalized volumetric cross-correlation (Supplementary Fig. 3D).

Fig. 2D shows a sequence of motion-corrected, background-subtracted (see Methods) SWIFT volumes taken during alternating periods of locomotion and non-locomotion of a CamKIIa::GCaMP6f-expressing mouse on the treadmill (Supplementary Movie 3). Note that parts of CA1, CA2, and CA3 are present in the 1.1 mm x 1.0 mm x 0.75 mm volumes shown (each is composed of 300 2.5um thick z-planes), and that individual cells are visible at depths 500 μm below the implant glass (white arrowhead in second volume; Fig. 2E shows z-slices through the indicated single time point volume). Imaging this curved structure across such depths in scattering tissue using only wide-field illumination is enabled by the maximum likelihood estimation method used for volume reconstruction (Broxton et al. 2013) (Methods), which can be thought of as yielding point estimates of each location in the presence of isotropic noise (e.g., that resulting from nearly isotropic scattering in turbid tissues).

Once motion corrected volumes are obtained, source extraction was performed on the resulting volumetric time series, resulting in the identified neurons shown in Fig. 2F,G. Fig. 2G shows 12 estimated calcium traces corresponding to the spatial filters of the numbered red neurons identified in Fig. 2F. Thresholded spike probability estimates (5% percentile) obtained using nonnegative deconvolution with a autoregressive model of the calcium transient (Vogelstein et al. 2010; Pnevmatikakis et al. 2016) was used to obtain the black dots below each trace, with each dot indicating a bin in which the cell is likely firing based on the model (Methods, scale bar 1 min).

## Volumetric imaging with cellular resolution during a contextual learning behavior

To leverage synchronous volumetric imaging across hippocampal subfields, we developed a virtual environment apparatus appropriate for contextual learning, a task in which the different subfields are thought to have distinct roles (McNaughton and Nadel 1990; Sekino et al. 1997; Chevaleyre and Siegelbaum 2010; Wintzer et al. 2014). The apparatus is similar to others reported previously (Harvey et al. 2009). Two video projectors presented a virtual environment onto a spherical screen placed 20 cm from the mouse and covering 220 degrees of the mouse’s visual field of view (Figure 3A; Methods). Head-restrained mice ran on an axially-constrained spherical treadmill, with forward or backward movement of the ball read out by an optical mouse (Dombeck et al. 2007). Automated delivery of odorants via an olfactometer, auditory cues by two speakers placed behind the mouse, and a small feeding tube placed immediately under the animal’s mouth allowed multisensory stimulation and training using water rewards in water-deprived animals (Fig 3A, Methods).

Standard video game development software (Unity3D) interfacing with custom Python code was used to present virtual environments with motor activity and licking behavior (measured via infrared movies of the mouse) as behavioral outputs (Methods). During training animals (n=3) ran through a series of two possible 120 cm long virtual “contexts”, pseudo-randomly ordered and with each context defined by a distinct combination of visual environment, auditory cue, and odorant (Figure 3B). One context was not visible from the other (animals ran through an opaque black doorway into the next context). Odorants were delivered for 1 second immediately at the start of the context (Stettler and Axel 2009) (Methods), and auditory cues were presented through the duration of the context starting immediately upon context entry (∼75 dB, see Methods). Of the two context types encountered by the animal, one was rewarded (and occurred in only ∼20% of all context presentations) while the other was not rewarded (∼80% of contexts; Fig. 3C top). During training, each animal received water rewards of the same magnitude (100 μL) at a 1 second delay after context entry. After 7-10 days, previously treadmill-naïve animals learned to run rapidly through the contexts to collect water rewards (Fig. 3C bottom), stopping to lick the reward tube after each reward delivery (top trace, bottom panel, Fig. 3C shows example mouse position relative to start of context; bottom trace, bottom panel shows smoothed licking rate). All mice successfully learned this task.

**Fig. 3.**
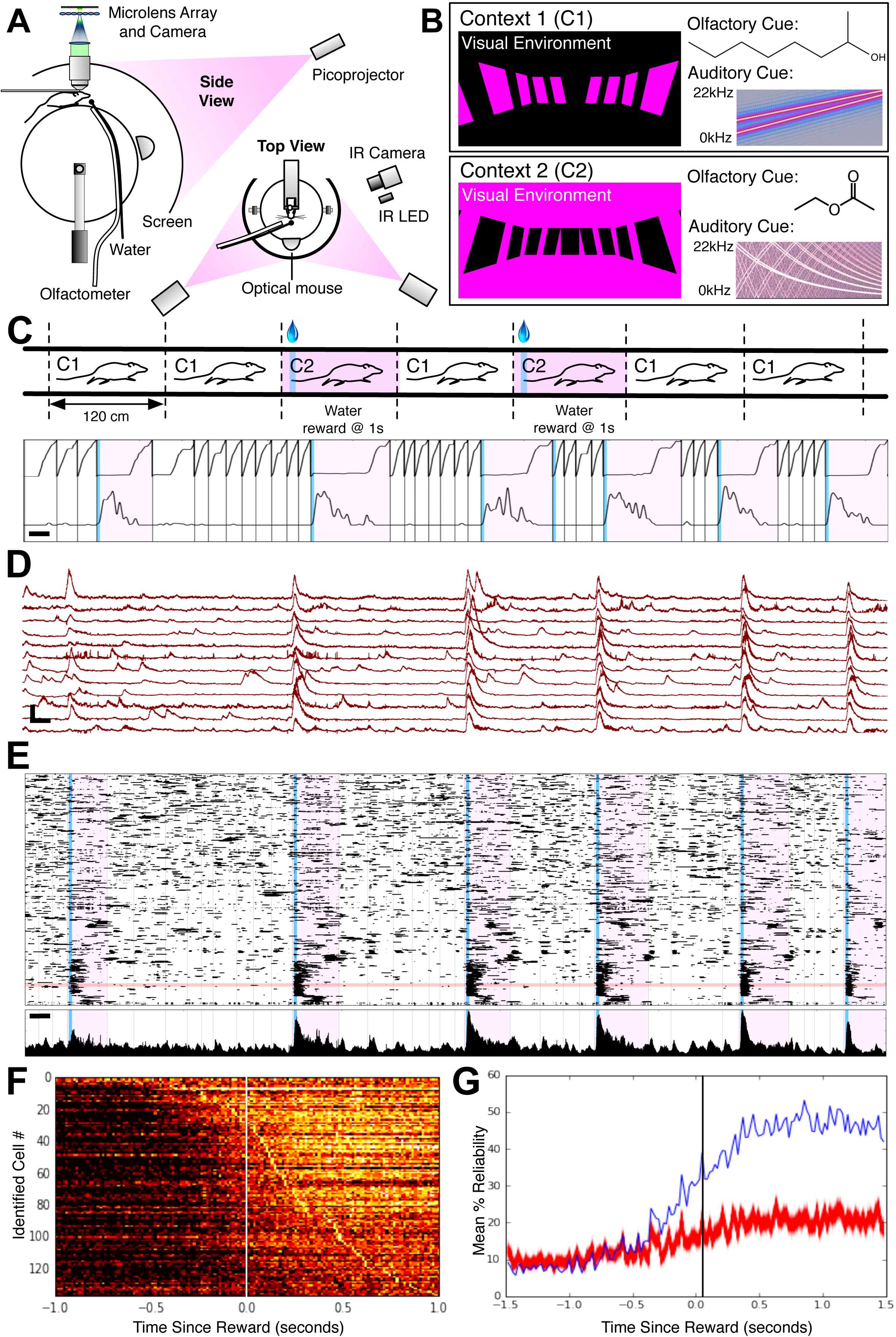
SWIFT volume imaging during a rewarded contextual navigation task. (**A**) (**B)** Mice are trained on a contextual reward task. (**C**) The mouse encounters two environments, C1 (white) and C2 (magenta underlay), which are defined by a distinct visual, olfactory, and auditory stimuli. Water-deprived mice receive no water reward in C1 trials (80% of trials), but they will receive a water reward 1 sec after entering C2 (20% of trials; blue line; scale bar 30 sec). (**D**) Twenty identified neurons that fire reliably in the rewarded context (highlighted in the horizontal red line in panel **E** below), and the corresponding activity traces (scale bar 400% dF/F, 30 sec). (**E**; top) Raster plots corresponding to spike likelihood traces (temporally-deconvolved and sorted by similarity, Methods); contexts and rewards indicated as in **C** (scale bar: 30 sec). (**E**; bottom), time histogram of the raster events. (**F**) Reliability of cells that fire within +/- 1 second of reward onset during >80% of trials. Cells are ordered vertically by consistency of firing. A bright pixel in the heatmap indicates the cell reliably fires during that time slice relative to administration of the reward. There is a temporal ordering to when cells fire reliably that starts prior to reward onset. (**G**) A permutation test (999 permutations, red lines) shows the reliability of cell firing in a time bin (blue line) to be significantly above chance (p<0.001, uncorrected), with divergence from the empirical null distribution beginning before reward onset.

SWIFT was used to locate cells across motion corrected volumes (Fig. 3D,E). Position in the context and licking behavior (Fig. 3C top and bottom traces, respectively) is registered to raster plots of calcium activity (Fig. 3E) for 1043 identified neurons (Methods) for one animal; and cells in the raster plot are dendrogram ordered by similarity of fluorescence traces (Methods). The activity traces and spatial locations of the 12 example neurons highlighted in the horizontal red band in Fig. 3E are shown in detail in Fig. 3D. We next looked at neurons that were active >80% of the time within a 2 second window centered around water reward onset (reward occurred reliably 1s after context entry; 139/1043 identified neurons responded >80%). Ordering these by the first time point at which their reliability exceeded the 80% threshold yielded a reliability heat map where each column represents a time point (relative to water reward onset) and each row corresponds to an identified neuron. Pixel color indicates the percentage of times the cell was active in that time bin across rewarded trials (Fig 3E). This plot reveals a ramping up of behavior beginning ∼50 ms before the reward is delivered, and persisting for at least 1s after reward onset in many cells. All three animals showed this similar pattern of reliability (n=3, 8.6% +- 0.44% SD of neurons showing >80% reliability; all starting after context entry and before reward onset). To determine if this activity was significantly different than expected by chance, we circularly permuted the time series within each trial (n=999 circular permutations) and show the empirically observed mean percent reliability in firing across neurons for each time bin (red) relative to the mean percent reliability in firing for circularly permuted time series. The observed trace clearly diverges from the empirical null significantly (p<0.001, uncorrected) ∼25ms prior to reward onset. All animals showed this divergence pattern over time, with initial significant divergence starting 25ms-50ms post context onset (n=3).

## Discussion

By integrating computational optics for volume reconstruction from two-dimensional camera images with high-dimensional statistics for motion correction and cell localization, SWIFT volume imaging enables fast, synchronous, cellular resolution recording and quantitative modeling of neural population activity across cortical layers and regions in scattering mammalian tissue. The frame rates of large-format scientific cameras and the kinetics of available fluorescent activity indicators therefore now represent the primary limiting factors for action-potential resolution imaging of large neural populations across brain regions. Increasing quality and speed of scientific cameras, and continued progress in the signal and speed of both calcium and voltage sensors will likely expand the spatiotemporal reach of the methods described here.

Computational imaging has spurred a revolution in photography (Giles 2009; Raskar 2009; M. Levoy 2010; Ehrenberg 2012), but so far has been relatively limited in its applications in biological light microscopy. Our findings highlight the recent promise of computational imaging to extend the reach of microscopy, using a combination of physical and statistical models, paired with new computational architectures for large data, to push the limits of classical imaging.

## Methods

### Light Field Microscopy

Imaging data were acquired using light field microscopes (Marc Levoy et al. 2006; Zhang and Levoy 2009; Broxton et al. 2013; Cohen et al. 2014). A standard commercial epifluorescence microscope (an Olympus BX51WI) was modified to be a light field microscope by placing a 125 μm pitch microlens array (RPC Photonics, see below for more details) at the intermediate image plane (i.e., the location of the camera sensor in a standard microscope). The back focal plane of the microlens array was then re-imaged via relay optics (see below) onto the sCMOS camera sensor (Andor Zyla, pixel size: 6.5μm). The microlens array and camera were placed on motorized linear stages (Thorlabs) for easy focusing adjustment.

We used microlens arrays with a square lenslet aperture (truncated, 100% fill factor) and a spherical lenslet surface profile. The f-number (*f*#) of the lenslets was chosen to match the magnification (M) and numerical aperture (NA) of the microscope objective via the formula: *f* # = *M*/(2*NA*). An f/10, 125 μm microlens array and a 10x 0.6 NA water-dipping objective (Olympus) were used in mouse in vivo/in vitro experiments. The relay optics between the microlens array and the camera sensor (Nikon NIKKOR) were chosen to demagnify the sensor pixels and increase the angular sampling rate in the light field. A 135:85 mm relay yielded 31 pixels per lenslet diameter.

Samples were illuminated evenly using LED epifluorescence illumination (Leica SP5 wide-field light engine or Lumencor Spectra X). All samples were imaged at wavelengths standard for GCaMP6f using fluorescence filters (Semrock filters: excitation 475/35 nm; dichroic 495 nm; emission 535/50 nm). Mouse in vitro volumes were recorded at 4 Hz or 25 Hz and mouse in vivo images at 50 Hz. We note that the maximum acquisition frame rate in our experiments is limited only by the frame rate of the camera. Although we started in mouse slice with longer integration times to ensure image data of sufficient quality to identify neurons in scattering tissue, we discovered that faster frame rates (up to 100 Hz in tests with GCaMP6f in vivo) were also effective. Volume imaging with frame rates up to 100 Hz and beyond are therefore possible using this technique, limited only by the speed and signal-to-noise ratio of the camera and calcium/voltage sensor signal to noise.

Light field images were recorded over a Camera Link 10-tap PCI interface onto a SSD RAID-0 disk array in a desktop workstation (Dell or Colfax Intl.). Images were recorded using our own custom image acquisition software as well as the open-source Micro-Manager image acquisition software (Edelstein et al. 2014). Dark frames and radiometric calibration frames were recorded before each experiment and used to correct for bias on the camera sensor and vignetting introduced by the microscope optics, respectively.

### Light field volume reconstruction/deconvolution

Volume data were reconstructed from raw light field frames using our light field three-dimensional deconvolution algorithm, which uses wave optics to model diffraction and other light transport phenomenon in the light field microscope. Three-dimensional reconstruction is formulated as an inverse problem that we solve using a GPU-accelerated iterative algorithm based on Richardson-Lucy iteration (Kralj et al. 2011; Gong et al. 2014). For more information on this algorithm and its implementation, we refer the reader to our previous study (Broxton et al. 2013).

Runtimes for our algorithm depend on the optical parameters used in a given experiment. In this study it took ∼2 minutes per camera frame for mouse in vivo and in vitro experiments. Between 3,000-27,000 frames are recorded for each experiment, so up to 1,000 machine hours are therefore required to process some data sets. To address this, we developed a distributed deconvolution data processing pipeline that runs on the Amazon Elastic Compute Cloud (Amazon EC2). Our Amazon EC2 cluster consists of 300 “g2.2xlarge” spot instances, each with a Kepler GK104 GPU (NVidia). We run four deconvolution jobs concurrently on each GPU, with jobs scheduled and managed using the Celery distributed task queuing system (http://www.celeryproject.org/). This GPU cluster has a theoretical peak computing power of 1,373 teraflops and is capable of processing all of the frames in a mouse in vivo dataset with 27,000 frames in 3 hours.

### Resolution limits

In light field microscopy, spatial resolution must be sacrificed in order to record different angles of light incident on the microscope’s image plane using different camera pixels (it is this angular information that permits scanless volumetric reconstruction). As a result, the resolution limit of a three dimensional light field reconstruction is higher than the Rayleigh limit encountered in wide field imaging. In general, spatial resolution gradually decreases at greater depths relative to the microscope’s native plane. We show in (Broxton et al. 2013) that this fall-off in lateral resolution is well approximated by a light field resolution criterion: *η*_*lf*_ = *d*/(0.94λ*M*|*z*|), where *d* is the pitch of the microlens array, λ is the emission wavelength of the sample, M is the magnification of the microscope objective, z is the distance relative to the microscope’s native image plane, and *η*_*lf*_ is the lateral spatial band limit (in cycles / mm) of a three-dimensional reconstruction for a given depth and set of optical parameters. This equation holds where *|z|* > *d*^2^/(2*M*^2^). Extended Data Fig. 1f shows this resolution limit plotted for various optical recipes used in this study. For a more in-depth discussion of these resolution limits, as well as experimental results validating them, we refer the reader to our previous study (Broxton et al. 2013).

### Sparse, structured matrix factorization for cell identification

Identification of candidate neurons (sources) required the development of a novel technique based on sparse and functional PCA (Allen 2013; Allen, Grosenick, and Taylor 2014). This technique (Allen 2013; Allen, Grosenick, and Taylor 2014) factors a matrix with columns representing time points and rows representing voxels into a sets of spatial coefficients and their corresponding time series where each set of coefficients is constrained to be both smooth and sparse—an assumption that has been shown to work well for locally smooth, volumetric neural data (Jenatton, Obozinski, and Bach 2009; Allen 2013; Grosenick et al. 2013; Pnevmatikakis et al. 2016). We augmented this technique with an additional connected component constraint that ensures that coefficients are non-negative, orthogonal, and restricted to a small region of space. This results in non-overlapping sets of coefficients each of which is connected in space. The augmented problem is non-convex, so we solved it heuristically using alternating minimization between the time series and spatial coefficients (Journée et al. 2010). We identified sources sequentially and initialized the optimization for each source using the highest scoring voxel that has not yet been included.

#### Initialization

Time series for each voxel were preprocessed to discount the effects of bleaching and baseline drift. A moving baseline (30 second window) was subtracted from each time series before it was standardized to have zero mean and unit norm. Due to the non-convex nature of the technique, it is sensitive to initialization. We initialized sources by choosing voxels that exhibit a large amount of positive deviation (peak) from a noisy baseline. Peaks were defined as the maximum difference between a smoothed time series with width *τ_on_*, and a smoothed time series with a width of 30 seconds. The noise level was computed as the mean absolute deviation between the raw time series and a smoothed time series (30 second window). The final initialization score was computed as the ratio of the peak to the noise level. This metric is highly sensitive to neurons firing a single spike throughout the entire experiment and is used to initialize, sort and screen candidate sources.

#### Screening

Candidate sources were screened to remove artifacts and other signals that were not likely to have originated from cell bodies. The screening criteria on sources were: (1) centroids could not be within 5 voxels (19.28 μm for mouse ex vivo and mouse in vivo datasets; the higher scoring source was chosen in the event of violation), (2) centroids could not be within 10 voxels of the lateral borders of the volume reconstruction, or within 5 voxels of the axial borders, (3) the number of nonzero voxels in a source must be between 2 and 1000, (4) the aspect ratio of the source (width / height of bounding box) must be between 0.5 and 2, and (5) the initialization score (defined above) must be greater than 2.

### Time series and raster plots

For time series plots, Delta F-over-F (dF/F) was computed as Δ*F* = (*x* – *F*_0_)/*F*_0_ where x was the time series, and *F*_0_ is its global mean. In raster plots, baseline and slow drift in baseline signal was removed from raw time series using a zero-phase temporal high pass filter (Butterworth, 1st order). Spiking events were then determined to be those that exceeded a 95 median absolute deviation (MAD) confidence interval. Finally, sources were hierarchically clustered (i.e. represented as a dendrogram) using the Pearson correlation coefficient a distance metric to compare pairs of time series.

### Mouse slice experiments

Female or male C57BL/6J mice (The Jackson Laboratory), aged 3–5 months at the time of imaging, were used for all experiments. They were housed on a 12-hour light/dark cycle in groups of 3–5 for acute slice imaging experiments and 2–3 for in vivo imaging experiments. The latter were also housed with a running wheel. All subjects were afforded ad libitum access to food and water, except during behavioral training as described below. All experimental protocols were approved by the Stanford University Institutional Animal Care and Use Committee.

Cloning of pAAV-CaMKIIa-GCaMP6f. The GCaMP6f was amplified from Addgene Plasmid #40755 using primers ccggatccgccaccatgggttctcatcatcatc and gataagcttgtcacttcgctgtcatcatttg, digested with BamHI and HindIII, and ligated to a AAV-CaMKIIa backbone cut with the same enzymes. Clones were verified by sequencing, and then the DNA was amplified and packaged as AAVdj at the Stanford Neuroscience Genomic Viral and Vector Core.

### Stereotactic viral injections

Viral injection was performed in mice anesthetized with 1.5-3.0% isoflurane and placed in a sterotaxic apparatus (Kopf Instruments). For in vitro imaging of the prefrontal cortex, mice were injected with 1000 nL of AAVdj-CKIIa-GCaMP6f (Stanford Vector Core) into the medial prefrontal cortex (coordinates relative to Bregma: AP +1.35 mm, ML –0.35 mm, DV –2.5 mm). For in vivo imaging of the hippocampus, mice were injected with 500nL of AAVdj-CKIIa-GCaMP6f in both CA1 (AP –1.5mm, ML –1.25mm, DV –1.6mm) and CA3 (AP –1.5mm, ML –1.75mm, DV –1.9mm), and injected with 250nL of AAV5-CKIIa-mCherry in the dentate gyrus (AP –1.5, ML –0.5, DV –2.15). Injections were performed using a 10 μL syringe and a 33 gauge beveled metal needle (Nanofil; WPI). Virus was extruded at a rate of 100nL/min using a syringe pump (UMP3; WPI) controlled by a Micro4 pump controller (WPI). Following virus injection, the incision was closed using tissue adhesive (Vetbond; Fisher) or sutures.

### Acute slice preparation for in vitro imaging

These experiments commenced 3–5 weeks after viral injections. Subjects were deeply anesthetized with 5% isoflurane and transcardially perfused with 20 mL of an ice-cold carbogenated solution of modified, protective artificial cerebrospinal fluid (composition: 92 mM NMDG, 25 mM glucose, 20 mM HEPES, 2.5 mM KCl, 1.2 mM NaH_2_PO_4_, 30 mM NaHCO_3_, 10mM MgSO_4_, 0.5 mM CaCl_2_, 5 mM sodium ascorbate, 2 mM thiourea, 3 mM sodium pyruvate; pH = 7.3–7.4; osmolarity = 295–305 mOsm). By substituting NMDG for sodium, perfusions with this solution may enhance slice viability by reducing excitotoxicity (Peça et al. 2011; Zhao et al. 2011). After perfusion, the brain was removed rapidly and cut into 350 um slices on a Leica vibratome using the same ice-cold NMDG-ACSF solution, and prefrontal slices were transferred immediately to a warm (32–34°C) recovery solution with the same composition. After 15 minutes of recovery, the slices were transferred to a modified HEPES-ACSF solution for 15 minutes (composition as above, except substituting 92 mM NaCl for NMDG), followed by a transfer to a second HEPES-ACSF solution to ensure complete washout of the NMDG. After 15 minutes the slices were transferred one more time to oxygenated ACSF solution and remained in this solution until imaging commenced (> 30 minutes later).

Imaging was conducted at a controlled 35°C in oxygenated ACSF (composition: 123 mM NaCl, 3 mM KCl, 11 mM glucose, 1.25 mM NaH_2_PO_4_, 26 mM NaHCO_3_, 1 mM MgCl_2_, 2 mM CaCl_2_) delivered to a Warner Instruments Ultra-Quiet Slice Chamber (RC-27LD) holding the slice through a in-line heater (Warner) at a constant rate of 8 mL/min through a peristaltic pump (Gilson). Standard “wide-field” illumination at 475/30 nm with a Lumencor Spectra X at less than 1 mW/mm^2^ was used for all imaging. Slice imaging datasets were acquired using 10X / 0.3 NA and 10X / 0.6 NA objectives (Olympus).

Hippocampal window implantation for in vivo imaging. These surgeries were performed after allowing at least one day of recovery after viral injections. Prior to the surgery, mice were injected with 80mg/6mg/kg of ketamine/xylazine intraperitoneally, and 5mg/kg carpofen subcutaneously. Mice were kept under 0.5-1.0% isoflurane throughout the surgery. First, a metal headplate was centered over the CA1 and CA3 injection sites and adhered to the skull using adhesive cement (Metabond; Parkell). A ∼3mm craniotomy was made in the center of the headplate opening using a trephine (Fisher). The cortex was then slowly aspirated using a 27 gauge blunt needle attached to a vacuum line while irrigating with lactated Ringer’s solution. Once a column of cortex had been removed, a 31 gauge blunt needle was used to peel away the white matter lying above the hippocampus. A glass implant was then lowered until the bottom coverslip rested against the hippocampus. The implant was constructed from 3.0mm diameter glass capillary tubes (Friedrich & Dimmock) custom cut to 1.5-1.75mm length, adhered on one end to a 3.0mm diameter coverslip of #0 thickness (Warner Instruments) using optical glue (Norland Products). The top of the implant extruding from the brain was then secured to the skull using adhesive cement.

### Virtual Environment

Experiments were performed using a custom-designed virtual environment setup, inspired by previous work (Harvey et al. 2009). A 200mm diameter styrofoam ball (Graham Sweet Studios) was fixed on a single rotational axis using a metal rod (6mm diameter cage assembly rods; Thorlabs) passing through the center axis of the ball and epoxied into place. The rod rested in two 90 deg post holders (Thorlabs), allowing free forwards and backwards rotation of the ball. Mice were head-fixed in place above the center of the ball using a headplate mount (Niell and Stryker 2010). Water rewards were delivered to mice through a small animal feeding tube (Cadence Science) connected to a 10mL syringe. A motorized syringe pump (Chemyx Inc.) delivered small water rewards triggered by a TTL pulse driven by custom control software (Python). Odorants were delivered through a multi-solenoid valve system (Parker Hannifin) connected to a pressurized air source using parameters from Stettler et al. (Stettler and Axel 2009). Auditory stimuli were uploaded into the virtual environment software and played over speakers (Logitech) located 40 cm symmetrically behind the mice. The mouse’s movements on the ball were recorded using an optical computer mouse (Logitech) that interfaced with the virtual environment software (Dombeck et al. 2007; Harvey et al. 2009). Virtual environments were designed in game development software Unity3d (unity3d.com). The virtual environment was displayed by back-projection onto projector screen fabric stretched over a clear acrylic hemisphere with a 14 inch diameter placed ∼20 cm in front of the center of the mouse. The screen encompasses ∼220 deg of the mouse’s field of view. The virtual environment was back-projected onto this screen using two laser-scanning projectors (Microvision), each projector covering one half of the screen. To create a flat image on the 3d screen, we warped the 2d image of the virtual environment using video manipulation software (Madmapper). The game engine allowed scripts written in Javscript or C# to trigger external events based on the mouse’s interactions with the virtual environment by communicating over a TCP socket to custom Python control software. A LabJackU6 (http://labjack.com/) was used both to time-lock virtual environment events and imaging frame times, and to send TTL pulses to deliver water rewards and odorants. The mouse’s licking behavior was recorded using a high-speed camera (Allied Guppy Pro; AVT) and 18mm lens (Edmund Optics), illuminated with an infrared LED (Thorlabs).

### Behavioral Task

Mice were water restricted for ∼5 days, or until reaching 80% of their starting weight, prior to the start of behavioral training. The behavioral paradigm consisted of two contexts, each a corridor of the same width (15 cm) and length (variable per mouse, see below). Each context differed in terms of the visual, olfactory, and auditory stimuli present. Context 1 was a black corridor with broad backwards-slanting magenta stripes on the walls. During Context 1 trials, a linear-sweeping tone (10 kHz to 25 kHz) was played, and a single puff of octanol was delivered at the start of the trial (odor delivery parameters were taken from Stettler et al. (Settler and Axel 2009). Context 1 was never associated with a water reward. Context 2 was a magenta corridor with broad forwards-slanting black stripes on the walls. During Context 2 trials, a logarithmic-sweeping tone (between 20 kHz and 22 kHz) was played, and a single puff of ethyl acetate was delivered at the start of the trial. Mice were given a water reward in Context 2. For each mouse, the lengths of the two contexts were set by adjusting the linear sensitivity of the optical mouse so that the non-rewarded Context 1 trials were completed in an average of 4-6 seconds. This was generally ∼120 cm. For a given mouse, the effective lengths of the two contexts were equal. Each training session consisted of 20 minutes, during which Context 1 and Context 2 trials were pseudo-randomly interleaved, with Context 1 trials occurring 80% of the time. Mice were trained on a task in which they were given water rewards 1s after entering Context 2.

---

**Supplementary Fig. 1 Comparison of confocal microscopy and light field microscopy. (A-E**) A 250 μm thick volume containing 2 μm fluorescent beads suspended in 2% low melting point agarose imaged with scanning confocal microscopy (**A,D**, 20x 0.5NA objective; scale bar 100 μm) and light field microscopy (**C,E**, same objective) (Marc Levoy et al. 2006; Broxton et al. 2013). **(B**) A raw light field image, where the pattern on the sensor generated by a microlens array encodes spatial and angular information about light emitted by the sources in the volume. Insets show different intensity patterns recorded for beads at (red) or below (blue) the native plane of focus of the microscope. (**C,E**) After reconstruction, the three-dimensional distribution of the beads can be distinguished in maximum intensity projections (**C**) and volume renderings (**E**). (**F**) Resolution criterion for the light field microscope showing the maximum resolvable spatial frequencies as a function of depth for several microscope objectives used in this study. For a fixed 125 μm pitch microlens array, a larger objective magnification results in better peak resolution but a more rapid fall-off and hence a diminished axial range over which good resolution can be achieved (Broxton et al. 2013).

**Supplementary Fig. 2. SWIFT volume imaging and validation in prefrontal cortex in vitro.** (**A**) Injection site for AAVdj-hSyn-GCaMP6f virus in mouse prefrontal cortex. (**B**) 864 active neurons identified across cortical layers 1, 2/3, 5, and 6 of an acute mouse cortical slice expressing genetically-encoded calcium indicator GCaMP6f in cingulate (Cg), prelimbic (PL), and infralimbic (IL) cortex using a novel sparse, structured machine learning method appropriate for massive data (Methods; scale bar 200 μm). (**C**) SWIFT fluorescence time series from twelve representative cells (those highlighted in red in **B**) (scale bar 580 %dF/F, 20 sec). (**D**) Fluorescence time series from 12 adjacent cells (highlighted in blue in **B**) centered on a randomly-chosen cell and showing no detectable cross-talk between adjacent identified cells (scale bar 580 %dF/F, 20 sec). (**E**) A volume reconstruction of a neuron patched and filled with Alexafluor 594 (purple). The red channel shows the identified cell spatial filter which agrees well with the filled cell image; black points indicate the centroids of the closest 20 identified sources (scale bar 300 μm). (**F**) Example voltage responses to depolarizing and hyperpolarizing current steps in two driven cells that were imaged at 25 Hz first using 2PSLM and then SWIFT (scale bar 50 mV; 250 ms). (**G**), signal-to-noise ratio (SNR) for 2PSLM (light/dark green correspond to cell 1/cell 2) and SWIFT (light/dark magenta correspond to cell 1/cell 2) resulting from 5 runs of the protocol shown in **F** for each imaging modality (Methods). (**H**) Correlations as a function of distance of nearby cells to the patched and driven cells in **F**; these show no consistent cross-contamination with distance in the SWIFT time series even at GCaMP6f-saturating levels of stimulation (although some cells at irregular distances show likely physiologically connections; nearby cell centroids for cell 1 are shown as black points in **E**).

**Supplementary Fig. 3. Structure of SWIFT analysis pipeline. (A)** Optical model for a light field microscope. An epifluorescence microscope can be converted into a light field microscope by placing a microlens array at the native image plane. Light from each position in the volume produces a unique intensity pattern that can be decoded by solving a large inverse problem, thereby reconstructing volumetric information from a single camera image. **(B)** Volumes are recovered using light field deconvolution (Broxton et al. 2013), which runs in parallel on 300 “g2.2xlarge” GPU-enabled Amazon spot instances. A single data set containing 30,000 raw light field images (10 minutes of data acquired at 50 Hz) can be processed in roughly 3 hours. **(C)** Volumes processed using a robust rigid-body motion correction algorithm (RASL) (Peng et al. 2012), and then sources and their time series are identified using a sparse, structured matrix factorization of the volumetric time-series. Time series are further temporally deconvolved and potentially thresholded to provide firing likelihoods or spike timing estimates (Vogelstein et al. 2010; Pnevmatikakis et al. 2016). **(D)** Time series extracted from a vertical slice (yellow line) exhibit high frequency artifacts from tissue motion (top) that remain present in cross-correlation based correction (middle) but not RASL-based motion correction (bottom).

## References

Abrahamsson, Sara, Jiji Chen, Bassam Hajj, Sjoerd Stallinga, Alexander Y. Katsov, Jan Wisniewski, Gaku Mizuguchi, et al. 2012. “Fast Multicolor 3D Imaging Using Aberration-Corrected Multifocus Microscopy.” Nature Methods, December. Nature Publishing Group, 1–6.

Ahrens, Misha B., Michael B. Orger, Drew N. Robson, Jennifer M. Li, and Philipp J. Keller. 2013. “Whole-Brain Functional Imaging at Cellular Resolution Using Light-Sheet Microscopy.” Nature Methods 10 (5). nature.com: 413–20.

Aimon, Sophie, Takeo Katsuki, Logan Grosenick, Michael Broxton, Karl Deisseroth, and Ralph J. Greenspan. 2015. “Activity Sources from Fast Large-Scale Brain Recordings in Adult Drosophila.” bioRxiv. doi:10.1101/033803.

Akemann, Walther, Hiroki Mutoh, Amélie Perron, Jean Rossier, and Thomas Knöpfel. 2010. “Imaging Brain Electric Signals with Genetically Targeted Voltage-Sensitive Fluorescent Proteins.” Nature Methods 7 (8). nature.com: 643–49.

Allen, Genevera I. 2013. “Sparse and Functional Principal Components Analysis.” arXiv [stat.ML]. arXiv. http://arxiv.org/abs/1309.2895.

Allen, Genevera I., Logan Grosenick, and Jonathan Taylor. 2014. “A Generalized Least-Square Matrix Decomposition.” Journal of the American Statistical Association 109 (505). Taylor & Francis: 145–59.

Barretto, Robert P. J., Bernhard Messerschmidt, and Mark J. Schnitzer. 2009. “In Vivo Fluorescence Imaging with High-Resolution Microlenses.” Nature Methods 6 (7): 511–12.

Bouchard, Matthew B., Venkatakaushik Voleti, César S. Mendes, Clay Lacefield, Wesley B. Grueber, Richard S. Mann, Randy M. Bruno, and Elizabeth M. C. Hillman. 2015. “Swept Confocally-Aligned Planar Excitation (SCAPE) Microscopy for High-Speed Volumetric Imaging of Behaving Organisms.” Nature Photonics 9 (2). Nature Publishing Group: 113–19.

Broxton, Michael, Logan Grosenick, Samuel Yang, Noy Cohen, Aaron Andalman, Karl Deisseroth, and Marc Levoy. 2013. “Wave Optics Theory and 3-D Deconvolution for the Light Field Microscope.” Optics Express 21 (21): 25418–39.

Chen, Tsai-Wen, Trevor J. Wardill, Yi Sun, Stefan R. Pulver, Sabine L. Renninger, Amy Baohan, Eric R. Schreiter, et al. 2013. “Ultrasensitive Fluorescent Proteins for Imaging Neuronal Activity.” Nature 499 (7458). nature.com: 295–300.

Chen, T. W., T. J. Wardill, Y. Sun, S. R. Pulver, S. L. Renninger, A. Baohan, E. R. Schreiter, et al. 2013. “Ultrasensitive Fluorescent Proteins for Imaging Neuronal Activity.” Nature 499: 295–300.

Chevaleyre, Vivien, and Steven A. Siegelbaum. 2010. “Strong CA2 Pyramidal Neuron Synapses Define a Powerful Disynaptic Cortico-Hippocampal Loop.” Neuron 66 (4): 560–72.

Cohen, Noy, Samuel Yang, Aaron Andalman, Michael Broxton, Logan Grosenick, Karl Deisseroth, Mark Horowitz, and Marc Levoy. 2014. “Enhancing the Performance of the Light Field Microscope Using Wavefront Coding.” Optics Express 22 (20). Optical Society of America: 24817–39.

Cossart, Rosa, Dmitriy Aronov, and Rafael Yuste. 2003. “Attractor Dynamics of Network UP States in the Neocortex.” Nature 423 (6937): 283–88.

Cotton, R. James, Emmanouil Froudarakis, Patrick Storer, Peter Saggau, and Andreas S. Tolias. 2013. “Three-Dimensional Mapping of Microcircuit Correlation Structure.” Frontiers in Neural Circuits 7 (October). ncbi.nlm.nih.gov: 151.

Deisseroth, Karl, and Mark J. Schnitzer. 2013. “Engineering Approaches to Illuminating Brain Structure and Dynamics.” Neuron 80 (3). Elsevier: 568–77.

Dombeck, Daniel A., Anton N. Khabbaz, Forrest Collman, Thomas L. Adelman, and David W. Tank. 2007. “Imaging Large-Scale Neural Activity with Cellular Resolution in Awake, Mobile Mice.” Neuron 56 (1): 43–57.

Duemani Reddy, Gaddum, Keith Kelleher, Rudy Fink, and Peter Saggau. 2008. “Three-Dimensional Random Access Multiphoton Microscopy for Functional Imaging of Neuronal Activity.” Nature Neuroscience 11 (6). nature.com: 713–20.

Edelstein, Arthur D., Mark A. Tsuchida, Nenad Amodaj, Henry Pinkard, Ronald D. Vale, and Nico Stuurman. 2014. “Advanced Methods of Microscope Control Using μManager Software.” Journal of Biological Methods 1 (2). doi:10.14440/jbm.2014.36.

Ehrenberg, Rachel. 2012. “The Digital Camera Revolution: Instead of Imitating Film Counterparts, New Technologies Work with Light in Creative Ways”. Science News 181 (2). John Wiley & Sons, Inc.: 22–25.

Freeman, Jeremy, Nikita Vladimirov, Takashi Kawashima, Yu Mu, Nicholas J. Sofroniew, Davis V. Bennett, Joshua Rosen, Chao-Tsung Yang, Loren L. Looger, and Misha B. Ahrens. 2014. “Mapping Brain Activity at Scale with Cluster Computing.” Nature Methods 11 (9): 941.50.

Giles, Jim. 2009. “Computational Cameras Perfect Your Photos for You. New Scientist, November 11. https://www.newscientist.com/article/mg20427346-700-computational-cameras-perfect-your-photos-for-you/.

Göbel, Werner, Björn M. Kampa, and Fritjof Helmchen. 2007. “Imaging Cellular Network Dynamics in Three Dimensions Using Fast 3D Laser Scanning.” Nature Methods 4 (1). nature.com: 73–79.

Gong, Yiyang, Mark J. Wagner, Jin Zhong Li, and Mark J. Schnitzer. 2014. “Imaging Neural Spiking in Brain Tissue Using FRET-Opsin Protein Voltage Sensors.” Nature Communications 5 (April): 3674.

Grosenick, Logan, Todd Anderson, and Stephen J. Smith. 2009. “Elastic Source Selection for in Vivo Imaging of Neuronal Ensembles. In Biomedical Imaging: From Nano to Macro, 2009.” ISBI’09. IEEE International Symposium on, 1263–66. IEEE.

Grosenick, Logan, Brad Klingenberg, Kiefer Katovich, Brian Knutson, and Jonathan E. Taylor. 2013. “Interpretable Whole-Brain Prediction Analysis with GraphNet.” NeuroImage 72(May). Elsevier: 304–21.

Harvey, Christopher D., Forrest Collman, Daniel A. Dombeck, and David W. Tank. 2009. “Intracellular Dynamics of Hippocampal Place Cells during Virtual Navigation.” Nature 461 (7266): 941–46.

Jenatton, Rodolphe, Guillaume Obozinski, and Francis Bach. 2009. “Structured Sparse Principal Component Analysis.” International Conference on Artificial Intelligence and Statistics, September.

Jin, Lei, Zhou Han, Jelena Platisa, Julian R. A. Wooltorton, Lawrence B. Cohen, and Vincent A. Pieribone. 2012. “Single Action Potentials and Subthreshold Electrical Events Imaged in Neurons with a Fluorescent Protein Voltage Probe.” Neuron 75 (5). Elsevier: 779–85.

Journée, Michel, Yurii Nesterov, Peter Richtárik, and Rodolphe Sepulchre. 2010. “Generalized Power Method for Sparse Principal Component Analysis.” Journal of Machine Learning Research: JMLR 11. JMLR.org: 517–53.

Kralj, Joel M., Daniel R. Hochbaum, Adam D. Douglass, and Adam E. Cohen. 2011. “Electrical Spiking in Escherichia Coli Probed with a Fluorescent Voltage-Indicating Protein.” Science 333 (6040): 345–48.

Levoy, M. 2010. “Experimental Platforms for Computational Photography.” IEEE Computer Graphics and Applications 30 (5): 81–87.

Levoy, Marc, Ren Ng, Andrew Adams, Matthew Footer, and Mark Horowitz. 2006. “Light Field Microscopy.” In ACM SIGGRAPH 2006 Papers, 924–34. SIGGRAPH ’06. New York, NY, USA: ACM.

Levoy, Marc, Zhengyun Zhang, and Ian McDowall. 2009. “Recording and Controlling the 4D Light Field in a Microscope Using Microlens Arrays.” Journal of Microscopy 235 (2). Wiley Online Library: 144–62.

McNaughton, Bruce L., and Lynn Nadel. 1990. “Hebb-Marr Networks and the Neurobiological Representation of Action in Space.” Neuroscience and Connectionist Theory. Erlbaum Hillsdale, NJ, 1–63.

Miyawaki, A., J. Llopis, R. Heim, J. M. McCaffery, J. A. Adams, M. Ikura, and R. Y. Tsien. 1997. “Fluorescent Indicators for Ca2+ Based on Green Fluorescent Proteins and Calmodulin.” Nature 388 (6645). nature.com: 882–87.

Niell, Cristopher M., and Michael P. Stryker. 2010. “Modulation of Visual Responses by Behavioral State in Mouse Visual Cortex.” Neuron 65 (4): 472–79.

Niesner, Raluca, Volker Andresen, Jens Neumann, Heinrich Spiecker, and Matthias Gunzer. 2007. “The Power of Single and Multibeam Two-Photon Microscopy for High-Resolution and High-Speed Deep Tissue and Intravital Imaging.” Biophysical Journal 93 (7). Elsevier: 2519–29.

Ntziachristos, Vasilis. 2010. “Going Deeper than Microscopy: The Optical Imaging Frontier in Biology.” Nature Methods 7 (8). Nature.com: 603–14.

Peça, João, Cátia Feliciano, Jonathan T. Ting, Wenting Wang, Michael F. Wells, Talaignair N. Venkatraman, Christopher D. Lascola, Zhanyan Fu, and Guoping Feng. 2011. “Shank3 Mutant Mice Display Autistic-like Behaviours and Striatal Dysfunction.” Nature 472 (7344): 437–42.

Peng, Yigang, Arvind Ganesh, John Wright, Wenli Xu, and Yi Ma. 2012. “RASL: Robust Alignment by Sparse and Low-Rank Decomposition for Linearly Correlated Images.” IEEE Transactions on Pattern Analysis and Machine Intelligence 34 (11): 2233–46.

Pnevmatikakis, Eftychios A., Daniel Soudry, Yuanjun Gao, Timothy A. Machado, Josh Merel, David Pfau, Thomas Reardon, et al. 2016. “Simultaneous Denoising, Deconvolution, and Demixing of Calcium Imaging Data.” Neuron 89 (2): 285–99.

Prevedel, Robert, Young-Gyu Yoon, Maximilian Hoffmann, Nikita Pak, Gordon Wetzstein, Saul Kato, Tina Schrödel, et al. 2014. “Simultaneous Whole-Animal 3D Imaging of Neuronal Activity Using Light-Field Microscopy.” Nature Methods 11 (7): 727–30.

Quirin, Sean, Jesse Jackson, Darcy S. Peterka, and Rafael Yuste. 2014. “Simultaneous Imaging of Neural Activity in Three Dimensions.” Frontiers in Neural Circuits 8 (April). ncbi.nlm.nih.gov: 29.

Raskar, Ramesh. 2009. “Computational Photography.” In Frontiers in Optics 2009/Laser Science XXV/Fall 2009 OSA Optics & Photonics Technical Digest, CTuA1. Optical Society of America.

Rosen, J., and G. Brooker. 2008. “Non-Scanning Motionless Fluorescence Three-Dimensional Holographic Microscopy.” Nature Photonics 2 (3). Nature Publishing Group: 190–95.

Sekino, Y., K. Obata, M. Tanifuji, M. Mizuno, and J. Murayama. 1997. “Delayed Signal Propagation via CA2 in Rat Hippocampal Slices Revealed by Optical Recording.” Journal of Neurophysiology 78 (3): 1662–68.

Stettler, Dan D., and Richard Axel. 2009. “Representations of Odor in the Piriform Cortex.” Neuron 63 (6): 854–64.

St-Pierre, François, Jesse D. Marshall, Ying Yang, Yiyang Gong, Mark J. Schnitzer, and Michael Z. Lin. 2014. “High-Fidelity Optical Reporting of Neuronal Electrical Activity with an Ultrafast Fluorescent Voltage Sensor.” Nature Neuroscience 17 (6). nature.com: 884–89.

Tian, Lin, S. Andrew Hires, Tianyi Mao, Daniel Huber, M. Eugenia Chiappe, Sreekanth H. Chalasani, Leopoldo Petreanu, et al. 2009. “Imaging Neural Activity in Worms, Flies and Mice with Improved GCaMP Calcium Indicators.” Nature Methods 6 (12). nature.com: 875–81.

Tomer, Raju, Khaled Khairy, Fernando Amat, and Philipp J. Keller. 2012. “Quantitative High-Speed Imaging of Entire Developing Embryos with Simultaneous Multiview Light-Sheet Microscopy.” Nature Methods 9 (7). nature.com: 755–63.

Tomer, Raju, Matthew Lovett-Barron, Isaac Kauvar, Aaron Andalman, Vanessa M. Burns, Sethuraman Sankaran, Logan Grosenick, Michael Broxton, Samuel Yang, and Karl Deisseroth. 2015. “SPED Light Sheet Microscopy: Fast Mapping of Biological System Structure and Function.” Cell 163 (7): 1796–1806.

Tsien, R. Y. 1980. “New Calcium Indicators and Buffers with High Selectivity against Magnesium and Protons: Design, Synthesis, and Properties of Prototype Structures.” Biochemistry 19 (11). ACS Publications: 2396–2404.

Vogelstein, Joshua T., Adam M. Packer, Timothy A. Machado, Tanya Sippy, Baktash Babadi, Rafael Yuste, and Liam Paninski. 2010. “Fast Nonnegative Deconvolution for Spike Train Inference from Population Calcium Imaging.” Journal of Neurophysiology 104 (6): 3691–3704.

Wintzer, Marie E., Roman Boehringer, Denis Polygalov, and Thomas J. McHugh. 2014. “The Hippocampal CA2 Ensemble Is Sensitive to Contextual Change.” The Journal of Neuroscience: The Official Journal of the Society for Neuroscience 34 (8): 3056–66.

Zaharia, Matei, Reynold S. Xin, Patrick Wendell, Tathagata Das, Michael Armbrust, Ankur Dave, Xiangrui Meng, et al. 2016. “Apache Spark: A Unified Engine for Big Data Processing.” Communications of the ACM 59 (11). New York, NY, USA: ACM: 56–65.

Zhang, Zhengyun, and M. Levoy. 2009. “Wigner Distributions and How They Relate to the Light Field.” In Computational Photography (ICCP), 2009 IEEE International Conference on, 1–10. IEEE.

Zhao, Shengli, Jonathan T. Ting, Hisham E. Atallah, Li Qiu, Jie Tan, Bernd Gloss, George J. Augustine, et al. 2011. “Cell Type-Specific Channelrhodopsin-2 Transgenic Mice for Optogenetic Dissection of Neural Circuitry Function.” Nature Methods 8 (9). Nature Publishing Group: 745–52.

